# Identification of Potential EGFR Inhibitors for Type 2 Diabetes and Pancreatic Cancer Treatment: A Computational Approach

**DOI:** 10.1101/2023.09.05.556125

**Authors:** Ricardo Romero

## Abstract

This study conducted a differential gene expression analysis in two independent studies of type 2 diabetes using pancreatic samples, specifically Langerhans cells. Through protein-protein interaction network analysis, the Epidermal Growth Factor Receptor (EGFR) emerged as the top hub gene among the upregulated genes in both studies. Furthermore, functional enrichment analysis revealed the involvement of EGFR in pancreatic cancer signaling pathways, indicating its potential role beyond diabetes.

To identify potential EGFR inhibitors, a virtual screening approach was employed using tyrosine kinase inhibitors. Two separate deep learning classification models were developed and trained on distinct sets of ligands, one for predicting bioactivity and the other for assessing toxicity. The ligands underwent rigorous filtering based on binding energies and absorption, distribution, metabolism, excretion, and toxicity (ADMET) properties.

This filtering process resulted in the identification of nine hits that demonstrated promising characteristics in terms of both bioactivity and druglikeness, making them viable candidates for further development as EGFR inhibitors.

To evaluate the performance of the identified hits, approved drugs including Afatinib, Almonertinib, Erlotinib, Gefitinib, and Osimertinib were used as controls. Notably, the finalist compounds consistently outperformed the control drugs across all evaluated parameters, indicating their superior potential as EGFR inhibitors.

This study provides valuable insights into the molecular mechanisms underlying type 2 diabetes by highlighting the significant role of EGFR and its potential association with pancreatic cancer signaling pathways. Moreover, the identified hits from the virtual screening represent promising candidates for further investigation and development of improved drugs targeting EGFR.

## 1 Introduction

Type 2 diabetes (T2D) is a complex and chronic metabolic disorder, caused by both genetic and environmental factors [1, 2, 3]. Its main characteristic is hyperglycemia, which results from impaired insulin secretion, insulin resistance, or both. It is estimated that over 400 million people worldwide are affected by T2D, and its prevalence continues to rise. Other characteristics include obesity, dyslipidemia, hypertension, and inflammation, which have been linked to the development of other complications such as cardiovascular disease, neuropathy, nephropathy, and retinopathy.

Several studies have shown that T2D is associated with reduced mass and/or reduced number of β-cells in the pancreatic islets [4, 5], which are clusters of endocrine cells located within the pancreas [6]. These cells are responsible for secreting hormones such as insulin, glucagon, and somatostatin, among others, as well as altered functionality of these cells [7], leading to insufficient insulin release and eventually hyperglycemia. Several genetic variants [8, 9, 10], identified through genome-wide association studies (GWAS), have been associated with the risk of developing T2D by influencing beta-cell function, mass, or response to various stimuli. Overall, the association between T2D and pancreatic islets highlights the importance of studying the underlying molecular mechanisms involved in the pathophysiology of disease onset and progression.

The management of T2D involves lifestyle changes such as diet and exercise, along with pharmacological prescriptions such as metformin, sulfonylureas, DPP4 inhibitors, etc., aiming to reduce blood sugar levels and improve insulin sensitivity [11, 12, 13]. However, these drugs do not stop or reverse the progression of the disease, focusing primarily on sugar control and symptom management. This highlights the need for better theoretical and experimental results that could eventually lead to the development of improved drugs for its treatment.

The purpose of the present study was to identify drug targets for the treatment of T2D and, based on this identification and using a computational approach, also identify compounds that could be repurposed or developed as inhibitory drugs for those targets.

As it turned out, the Epidermal Growth Factor Receptor (EGFR) resulted as the primary target for T2D according to the differential gene expression analyses performed. EGFR is a transmembrane protein also known as ErbB1 or HER1, which plays a critical role in cellular signaling pathways. It belongs to the ErbB family of tyrosine kinase receptors and is expressed on the surface of various cells, including epithelial cells. Activation of EGFR initiates a cascade of intracellular events that lead to cell proliferation, survival, differentiation, and migration. The main features of the human EGFR include its extracellular domain, responsible for recognizing growth factors such as epidermal growth factor (EGF) and transforming growth factor-alpha (TGF-α). Upon binding to its ligand, EGFR undergoes conformational changes that result in dimerization and auto-phosphorylation of specific tyrosine residues within its cytoplasmic kinase domain. This phosphorylation activates downstream signaling pathways involved in the regulation of cell growth, tissue development, wound healing, and maintenance of epithelial tissues. Dysregulation or mutations in EGFR have been associated with various cancers [14], making it an important target for therapeutic interventions, such as targeted therapies and small molecule inhibitors. Although the relation between EGFR and diabetes is not conclusive, there are already studies pointing to their association, such as the proposal of EGFR as a therapeutic target for diabetic kidney disease and insulin resistance [15, 16].

Several FDA-approved drugs [17], specifically designed to inhibit EGFR signaling, have proven to be effective treatments for certain types of cancers, including non-small cell lung cancer (NSCLC), colorectal cancer, and head and neck squamous cell carcinomas. These include Gefitinib (Iressa), which is an EGFR tyrosine kinase inhibitor approved for the treatment of NSCLC. Erlotinib (Tarceva) is another EGFR tyrosine kinase inhibitor indicated for NSCLC and pancreatic cancer. Afatinib (Gilotrif) is an irreversible tyrosine kinase inhibitor targeting both EGFR and HER2, used in NSCLC treatment with specific mutations. Osimertinib (Tagrisso) is a drug approved for advanced or metastatic NSCLC patients who have developed resistance to other EGFR inhibitors due to a T790M mutation. Cetuximab (Erbitux) is a monoclonal antibody targeting EGFR, used in colorectal cancer, head and neck cancer, and some types of lung cancer. These targeted therapies can provide personalized treatment options by selectively blocking EGFR activity or its downstream signaling pathways, leading to improved clinical outcomes for patients with EGFR-driven malignancies.

The identification of EGFR as a T2D target was obtained through differential gene expression analysis, interaction networks and hub genes identification based on two T2D studies with pancreatic islet samples. Subsequently, virtual screening was performed using deep learning models to predict bioactivity and toxicity, followed by docking. Finally, an ADMET filter was also implemented to obtain the final hits with the best characteristics to proceed to the in vitro phase.

## 2 Methods and materials

A flowchart depicting the steps followed in this study is shown in Figure 1.

**Figure 1.**
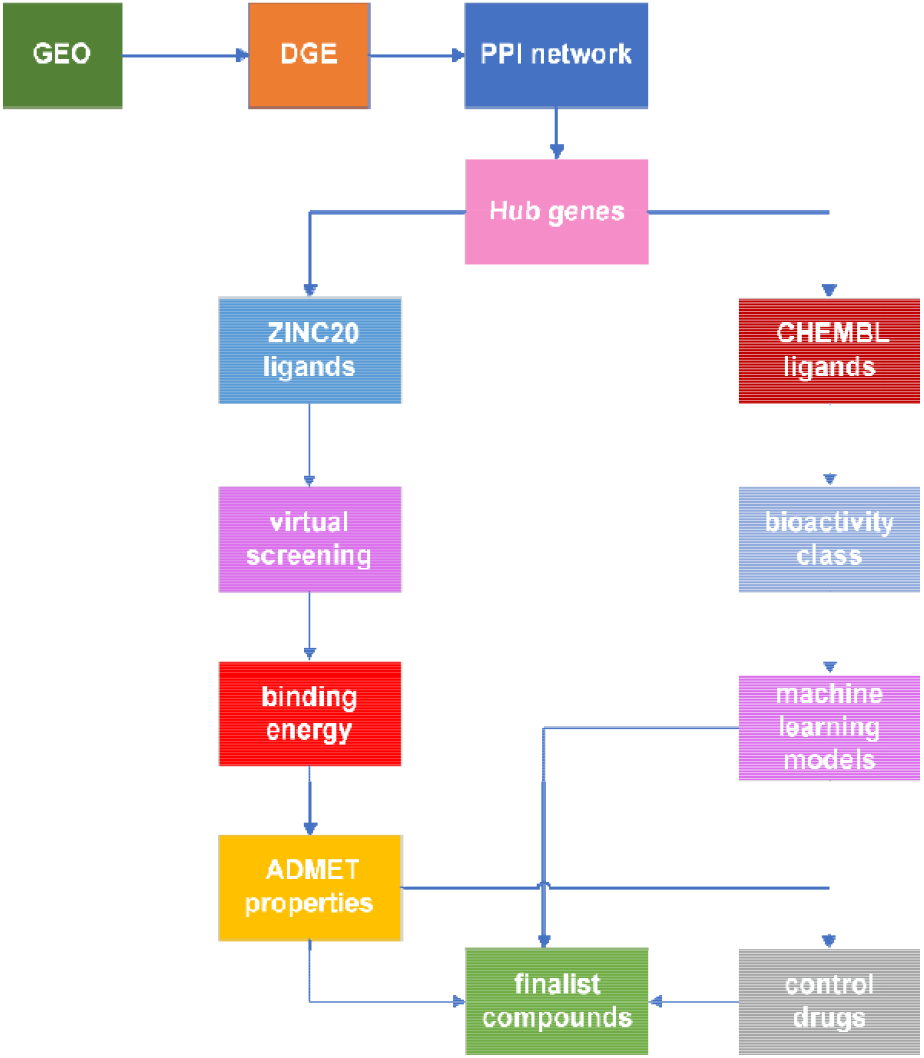
Flowchart of the procedure followed in the present study. The data is obtained from the Gene Expression Omnibus (GEO), and an analysis of differential gene expression (DGE) is performed. The protein-protein interaction (PPI) network is derived, and hub genes are identified. Ligands are obtained from ZINC20, and virtual screening is conducted. The results are filtered based on binding energy and ADMET properties. Simultaneously, machine learning models are trained to further filter compounds based on their bioactivity. The final compounds are compared with control drugs.

### 2.1 Data Acquisition and Analysis

Two studies, GSE25724 and GSE196797, were obtained from the Gene Expression Omnibus (https://www.ncbi.nlm.nih.gov/geo) database for the present analysis. These studies were selected based on the following criteria:

⁳ The studies focused on type 2 diabetes using pancreatic islet samples.
⁳ The datasets provided a clear distinction between type 2 diabetes (T2D) and non-T2D (control) samples.
⁳ The gene expression analysis yielded differentially expressed genes suitable for constructing a protein-protein interaction (PPI) rich network.

The purpose of the last requirement will be clarified in the results section. The GSE25724 study data was analyzed online using the GEO2R tool, a differential gene expression analysis tool provided by NCBI for microarray data. The analysis compared 6 T2D samples versus 7 non-T2D samples. Data was normalized and adjusted p-values were calculated using the Benjamini and Hochberg method. Logarithmic transformation and precise weights from the R limma package were also applied.

The significance level was 0.05 with a log2 fold change (log2FC) threshold of zero. The results were filtered by selecting genes with log2FC > 0 (since only overexpressed genes were of interest) and adjusted p-value < 0.05. The final data was then loaded into the STRING server (https://string-db.org) to obtain the PPI network, which was then imported into the Cytoscape software [18] to obtain the hub genes by means of the CytoHubba plugin [19]. The top ten hub were genes identified based on their degree of interaction.

For the GSE196797 study, which involved miRNA high-throughput sequencing, the data was analyzed in R Studio using the DESeq2 package [20] available from the Bioconductor resource center [21], since it was not available for analysis using the GEO2R tool. Normalized gene count data with 2 T2D samples versus 6 non-T2D samples were used. Genes with a p-value < 0.05 and log2FC > 0 were selected from the differential expression analysis. Here, the standard p value was used rather than the adjusted since it was not possible to obtain the latter for all genes. The PPI network and top ten hub genes were obtained following the same procedure as the previous study. The datasets and the R script used for the analysis are available in the supplementary materials.

### 2.2 Enrichment Analysis

Due to their richer PPI network, the top ten hub genes from the GSE25724 study were selected for enrichment analysis, which was performed in the Enrichr database [22, 23], with the hub genes compared against the KEGG 2021 Human signaling pathways database [24].

A transcriptomic analysis was performed using the GeneTrail 3.2 database [25], with both the hub genes and the study’s accession number. This analysis incorporated data from the Wiki Pathways 2021 Human database [26] and Gene Ontology (GO)[27, 28], and the results were identical in both cases, which served as an independent validation of the hub genes found.

To explore the involvement of miRNAs in diabetes and pancreatic neoplasms, groups of miRNAs associated with the hub genes were obtained from the miRNet database [29, 30]. This was accomplished by analyzing all the miRNAs linked to the hub genes using the miRNA disease database and performing a hypergeometric test.

### 2.3 Deep learning models

A deep learning model pipeline was implemented to filter the results from virtual screening. The first filter consisted of retaining only active compounds according to a multilayer perceptron (MLP) classification model trained on different data, while the second filter was also based on an MLP model, trained on the tox21 dataset to predict toxicity [31], this time requiring that the compounds passed all the twelve toxicity assays presented in the dataset.

To train the first model, data was downloaded and curated from the ChEMBL database [32]. It consisted of compounds targeting the human tyrosine kinases EGFR (CHEMBL203), VEGFR (CHEMBL1868), PDGFR (CHEMBL2007), FGFR (CHEMBL3650), ABL (CHEMBL1862) and SRC (CHEMBL267). The data was processed to remove zero and missing IC50 values, as well as duplicate and missing canonical SMILES.

The compounds were then classified as follows: compounds with an IC50 value ≤ 1000 nm were labeled active, those with an IC50 value ≥ 10000 nm were labeled inactive, and the remaining ones were classified as intermediate. The IC50 values were normalized to a cut-off value of 10^9 nm (since they were all given in nanomoles) and converted into pIC50 values to account for the great value range. Lipinski descriptors were also calculated, and a Mann-Whitney statistical test applied to verify that the active and inactive classes belonged to different distributions. The curated data as well as the code used to train the model are available in the Kagle database [33]. The model was trained using the pIC50 values in binary form as the target variable: 1 for pIC50 ≥ 6 (active) and 0 for the rest (non-active, including inactive and intermediate). The independent variables were the molecular fingerprints calculated using the Python package RdKit.

For the second model, curated data from the original tox21 dataset was downloaded and used to train it [34]. This time 1 indicated a positive result and 0 a null result for each of the twelve toxicity assays which comprised the target variables. Once the models were trained, they were validated on the testing sets and then applied to results from the virtual screening. The Python code used in this study is available in the supplementary materials.

### 2.3 Virtual Screening Procedure

Among the available PDB structures, entry 4HJO [35] was chosen as it consisted of the EGFR tyrosine kinase domain in complex with erlotinib, with the latter being a known EGFR inhibitor selected as one of the controls for the present study. The complex also served the purpose of identifying the molecule active site (Figure 2). Prior to docking, the structure was stripped of the ligand and water molecules using USCF Chimera [36], and energy minimization was performed using Swiss PDB-viewer 4.10 [37]. Hydrogens and Gasteiger charges were also assigned to the structure using Chimera.

**Figure 2.**
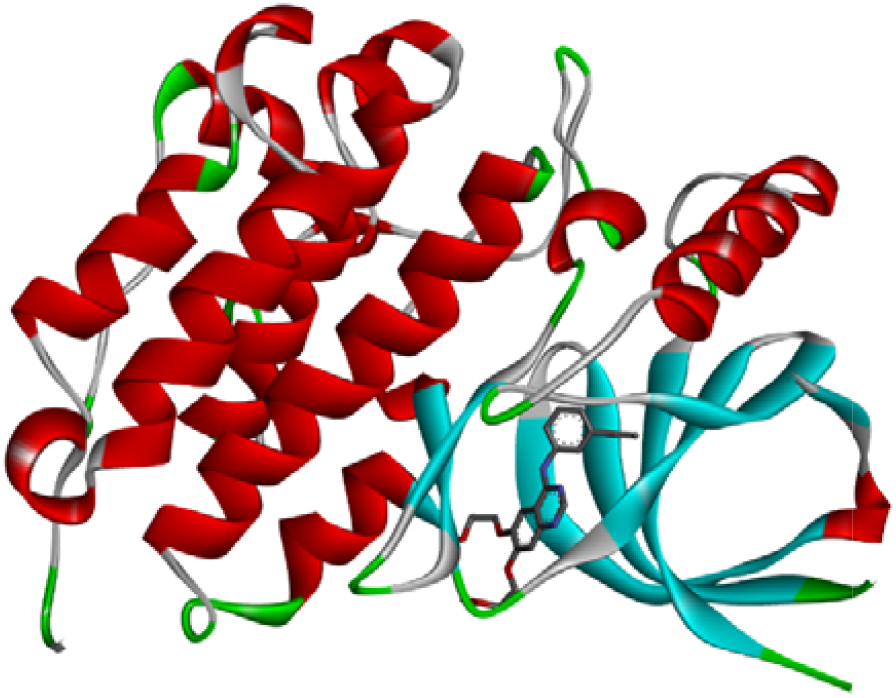
Structure of the Epidermal Growth Factor Receptor (EGFR) tyrosine kinase domain, PDB 4HJO, used for molecular docking. The picture shows the inactive molecule structure in complex with erlotinib in the active site of the tyrosine kinase domain. The 4HJO structure was obtained through X ray diffraction.

For virtual screening, 100 ligands targeting the EGFR protein, within the subset of predicted compounds, were downloaded from the ZINC20 database [38] and 36,324 compounds were downloaded from ChemDiv’s Protein Kinases Inhibitors Library [39]. The ligands were compared against the ones obtained from ChEMBL and duplicates eliminated before virtual screening. The compounds were then filtered by the deep learning models and proceeded to docking.

Energy minimization was carried out using the Amber force field with 5000 steps in the Open Babel software [40] and the docking procedure was performed with Autodock Vina [41, 42], using coordinates centered at (28.301, 2.272, 3.051) and a grid size of (20, 30, 20) to cover the entire active site of the tyrosine kinase domain, as determined by the complex with erlotinib. The exhaustiveness value was set to 30.

### 2.4 ADMET filtering

Following docking a third filter was implemented via the Swiss ADME database [43], with compounds required to pass the druglikeness rules of Lipinski [44], Ghose [45], Veber [46], Egan [47], and Muegge [48], as well as the structure Brenk [49] and PAINS alerts [50].

To further assess potential toxicological risks, the OSIRIS Property Explorer and Data Warrior programs were employed for a final filter, incorporating results for mutagenicity, tumorigenicity, irritancy, and reproductive effects. Compounds that passed these filters were recorded, along with their respective solubility, druglikeness score, and drug score values given by Osiris property explorer and the druglikeness score provided by Data Warrior. Detailed parameters calculations can be found on the programs’ websites.

To support the analysis, control drugs Afatinib, Almonertinib, Erlotinib, Gefitinib, and Osimertinib were included. These drugs were chosen based on their superior characteristics among each generation of approved EGFR inhibitors [17]. Their three-dimensional structures were downloaded from PubChem and optimized for docking using Avogadro [51], with the Generalized Amber Force Field (GAFF) with 5000 steps.

Using binding energy values from docking, along with data from the toxicity software, a simple average score was calculated using the formula: Score = 1/5(solubility + druglikeness (Osiris)+ druglikeness (DW) + Drug score + |binding energy|). Solubility was left without taking the absolute value because, although it is negative, less negativity indicates better solubility, unlike binding energy.

## 3 Results

The study yielded several significant findings. Firstly, nine compounds with suitable characteristics for potential EGFR inhibitors, eight from the ChemDiv database and one from ZINC, were identified and classified. Tables 1 and 2 provide detailed information about these compounds, along with the control drugs used for comparison. Canonical SMILES of the finalists’ compounds are provided in Table 4.

**Table 1.** Compounds properties. The binding energy from docking is given in kcal/mol. Solubility, Druglikeness (OPE), and Drug Score are the scores obtained from OSIRIS Property Explorer. Druglikeness (DW) is the score obtained by Data Warrior. The final score is calculated using the formula 1/5(solubility + druglikeness (OPE) + druglikeness (DW) + drug score + |binding energy|). The pIC50 column contains the numerical values predicted by the machine learning regression model, and the corresponding Class column presents the classification explained in section 2.5 (after conversion to IC50 values). All compounds were classified as active by the deep neural network bioactivity model.

| Compound | Binding Energy | Solubility | Druglikeness (OPE) | Drug score | Druglikeness (DW) | Final score |
| --- | --- | --- | --- | --- | --- | --- |
| E626-0366 | -9.2598 | -2.8 | 7.24 | 0.76 | 7.8474 | 4.46 |
| E626-0262 | -9.1277 | -3.59 | 6.85 | 0.66 | 6.54 | 3.92 |
| G430-1362 | -9.483 | -5.03 | 5.61 | 0.61 | 6.7329 | 3.48 |
| G430-1280 | -9.2518 | -5.4 | 7.29 | 0.56 | 5.4913 | 3.44 |
| E844-1573 | -9.2875 | -5.39 | 5.93 | 0.56 | 6.1287 | 3.30 |
| K405-4307 | -9.5103 | -5.36 | 3.27 | 0.58 | 7.1067 | 3.02 |
| J106-0699 | -9.1609 | -4.96 | 4.95 | 0.64 | 4.4441 | 2.85 |
| G430-1374 | -10.5569 | -5.35 | 3.45 | 0.56 | 5.3929 | 2.92 |
| ZINC6260 | -9.1317 | -4.43 | 0.36 | 0.65 | 1.8931 | 1.51 |
| Afatinib | -8.8336 | -5.48 | -4.11 | 0.58 | -0.79682 | -0.26 |
| Erlotinib | -8.0544 | -3.53 | -6.73 | 0.38 | -5.9718 | -1.56 |
| Gefitinib | -8.5039 | -5.06 | -2.62 | 0.28 | 0.47937 | 0.32 |
| Osimertinib | -8.6615 | -4.45 | -4.58 | 0.09 | -5.4185 | -1.14 |
| Almonertinib | -8.6505 | -4.46 | -4.09 | 0.08 | -4.9192 | -0.95 |

**Table 2.** Druglikeness and toxicity properties. Molecular weight (MW) in g/mol, topological surface area (TPSA) in Å^2^, brain blood barrier permeability (BBB). The Lipinski (L), Ghose (G), Veber (V), Egan (E) and Muegge (M) columns indicate the number of violations of the respective rules. The columns PAINS (Pan Assays Interference Structures) and Brenk indicate the number of alerts of the corresponding conditions. The columns Mutagenic, Tumorigenic, Reproductive effective and Irritant display the corresponding risks according to Data Warrior. All the compounds pass the 12 assays of the tox21 filter.

| Compound | MW | TPSA | BBB | L | G | V | E | M | PAINS | Brenk | Mutagenic | Tumorigenic | Reproductive effective | Irritant |
| --- | --- | --- | --- | --- | --- | --- | --- | --- | --- | --- | --- | --- | --- | --- |
| E626-0366 | 432.54 | 103.66 | No | 0 | 0 | 0 | 0 | 0 | 0 | 0 | none | none | none | none |
| E626-0262 | 465.54 | 100 | No | 0 | 0 | 0 | 0 | 0 | 0 | 0 | none | none | none | none |
| G430-1362 | 383.49 | 36.67 | Yes | 0 | 0 | 0 | 0 | 0 | 0 | 0 | none | none | none | none |
| G430-1280 | 407.87 | 36.67 | Yes | 0 | 0 | 0 | 0 | 0 | 0 | 0 | none | none | none | none |
| E844-1573 | 422.48 | 68.92 | Yes | 0 | 0 | 0 | 0 | 0 | 0 | 0 | none | none | none | none |
| K405-4307 | 384.48 | 50.08 | Yes | 0 | 0 | 0 | 0 | 0 | 0 | 0 | none | none | none | none |
| J106-0699 | 414.26 | 90.98 | No | 0 | 0 | 0 | 0 | 0 | 0 | 0 | none | none | none | none |
| G430-1374 | 401.48 | 36.67 | Yes | 0 | 0 | 0 | 0 | 0 | 0 | 0 | none | none | none | none |
| ZINC6260 | 289.33 | 55.63 | Yes | 0 | 0 | 0 | 0 | 0 | 0 | 0 | none | none | none | none |
| Afatinib | 485.94 | 88.61 | No | 0 | 1 | 0 | 0 | 0 | 0 | 1 | none | none | none | none |
| Erlotinib | 393.44 | 74.73 | Yes | 0 | 0 | 0 | 0 | 0 | 0 | 1 | none | none | none | none |
| Gefitinib | 446.9 | 68.74 | Yes | 0 | 0 | 0 | 0 | 0 | 0 | 0 | none | none | none | none |
| Osimertinib | 499.61 | 87.55 | No | 0 | 2 | 1 | 0 | 0 | 0 | 1 | high | low | low | low |
| Almonertinib | 525.64 | 87.55 | No | 1 | 3 | 1 | 0 | 0 | 0 | 1 | high | low | low | low |

Results from gene expression analysis, IPP networks and hub genes, for both studies used, are shown in figures 3 to 6. Notably, the EGFR hub gene exhibited the highest degree of interaction in both cases. Molecular docking results for the two highest-scoring compounds and from the ZINC ligand are summarized in Table 3. Figures 7 to 9 illustrate the complexes derived from these docking analyses.

**Table 3.**
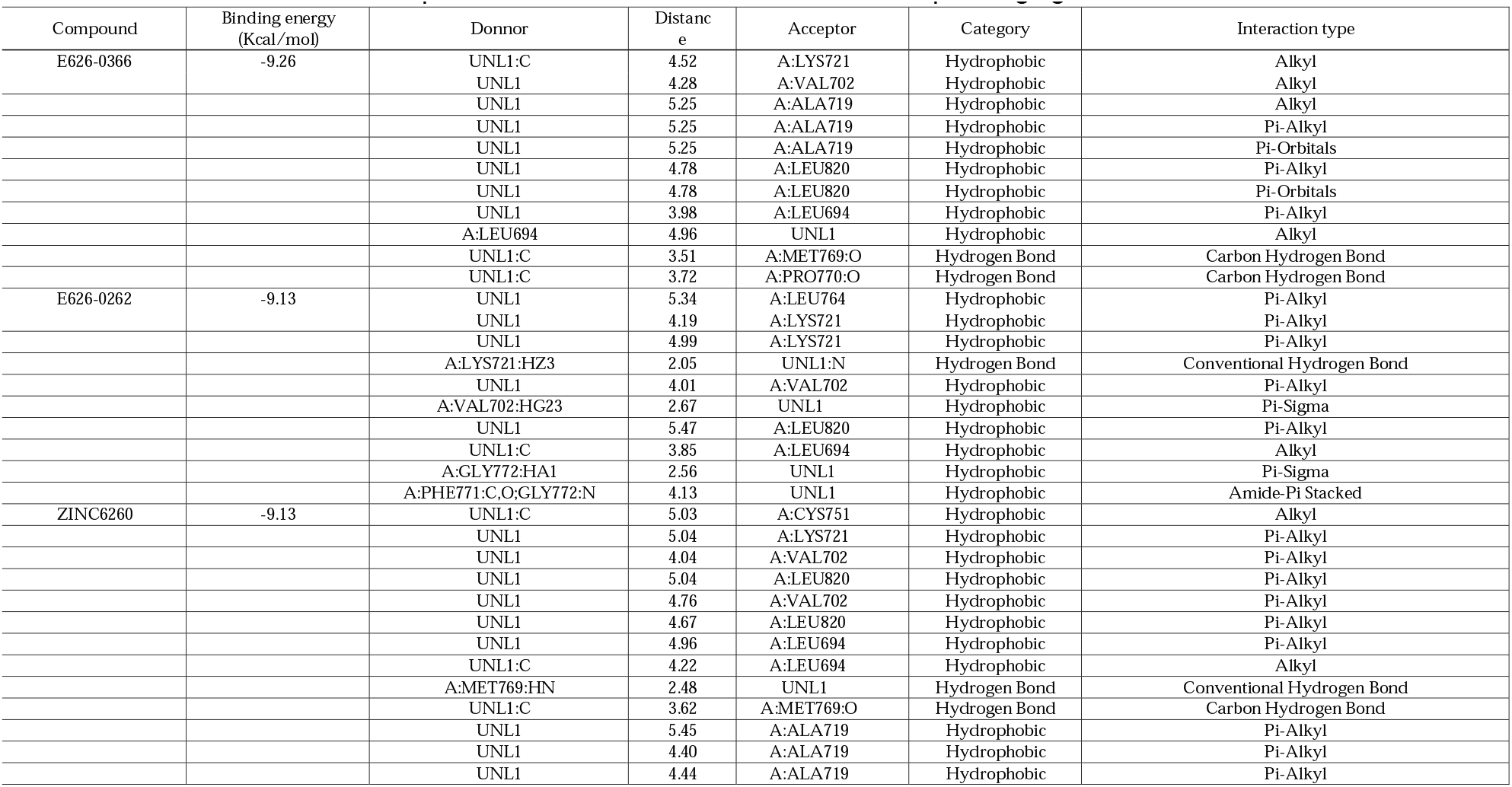
Binding energy and type of molecular interaction for the three finalist compounds E626-0366, E626-0262 and ZINC6260.The distance is reported in Å^2^ and UNL1 refers to the corresponding ligand in each case.

**Figure 3.**
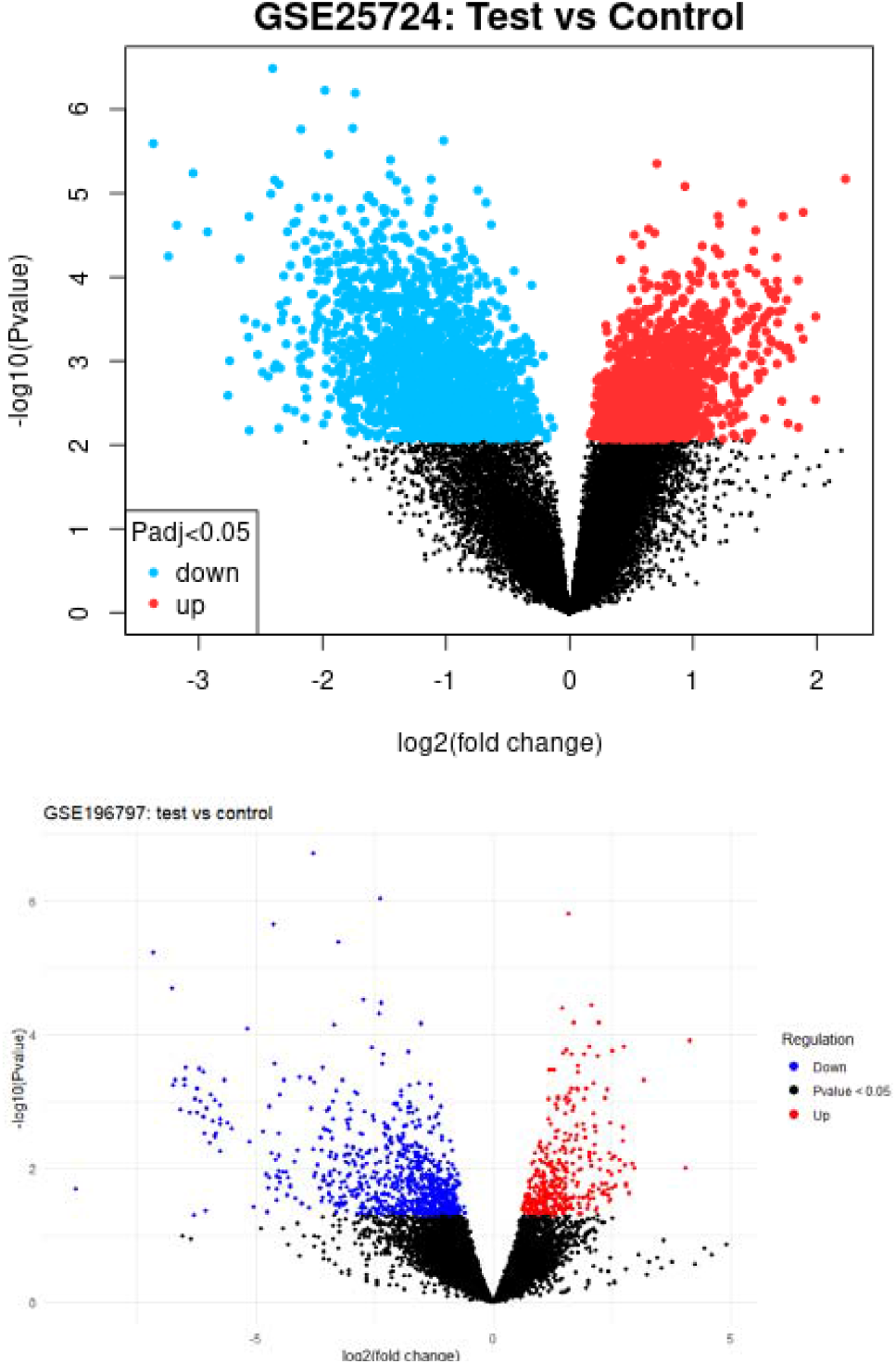
Volcano plots for the differentially expressed genes in both studies.

**Figure 4.**
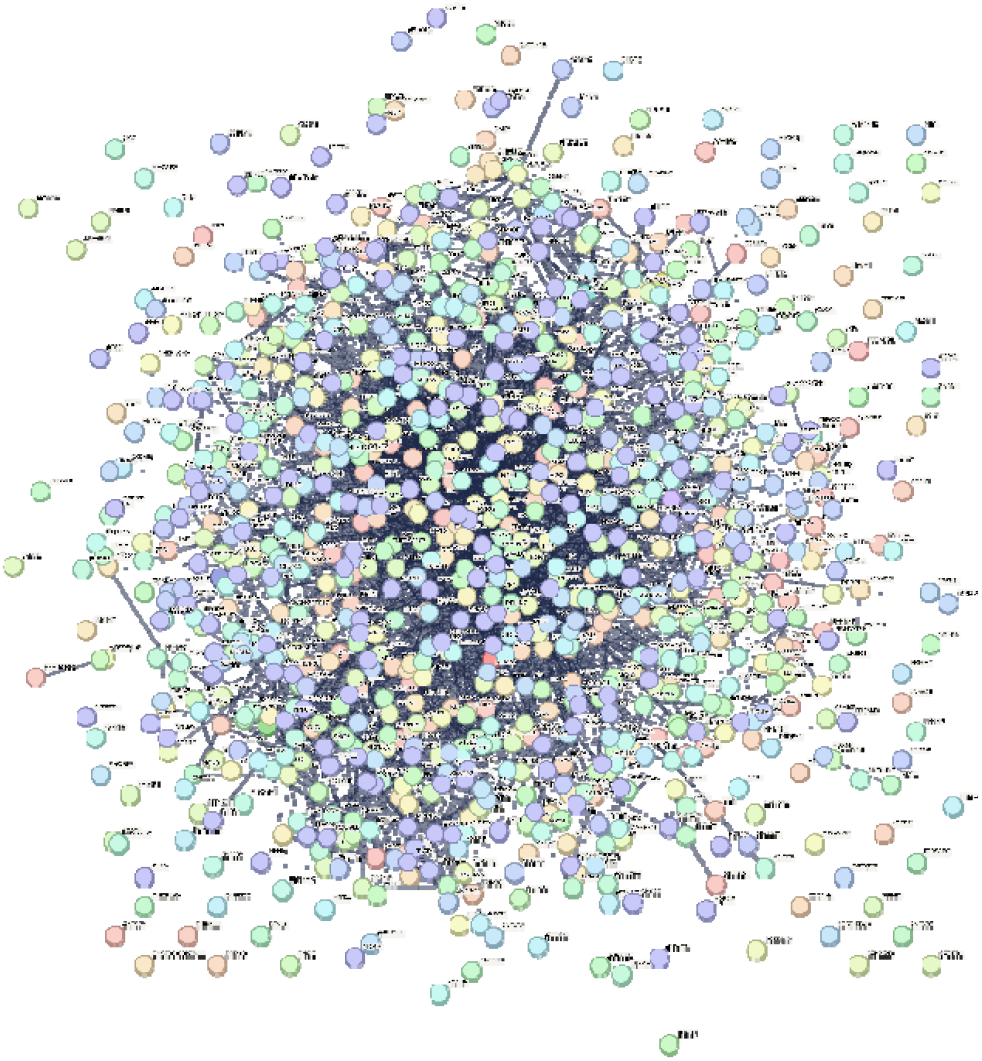
Protein-Protein Interaction (PPI) network for the up-regulated genes in the GSE25724 study. It consists of 998 nodes and 5036 edges, with an average node degree of 10.1 and an average local clustering coefficient of 0.338. The expected number of edges is 4699, with a PPI enrichment p-value of 5.99e-07.

**Figure 5.**
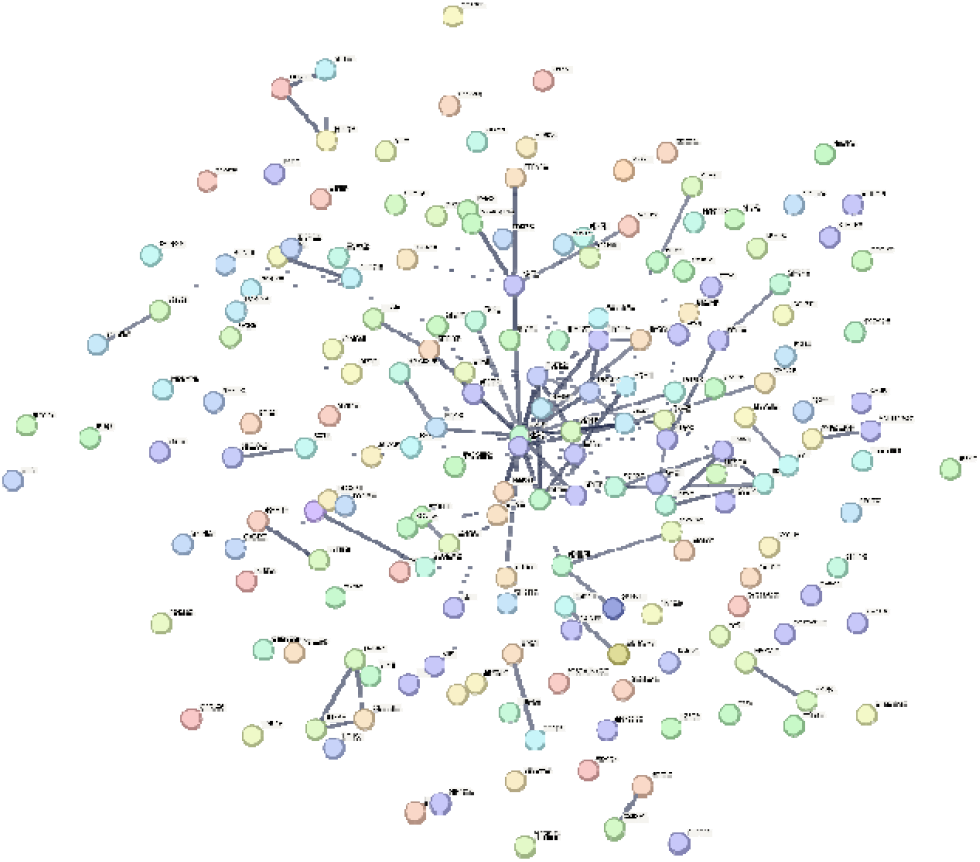
PPI network for up-regulated genes in study GSE196797. It consists of 298 nodes and 314 edges, with an average node degree of 2.11 and an average local clustering coefficient of 0.314. The expected number of edges is 229 with a PPI enrichment p-value of 6.37e-08.

**Figure 6.**
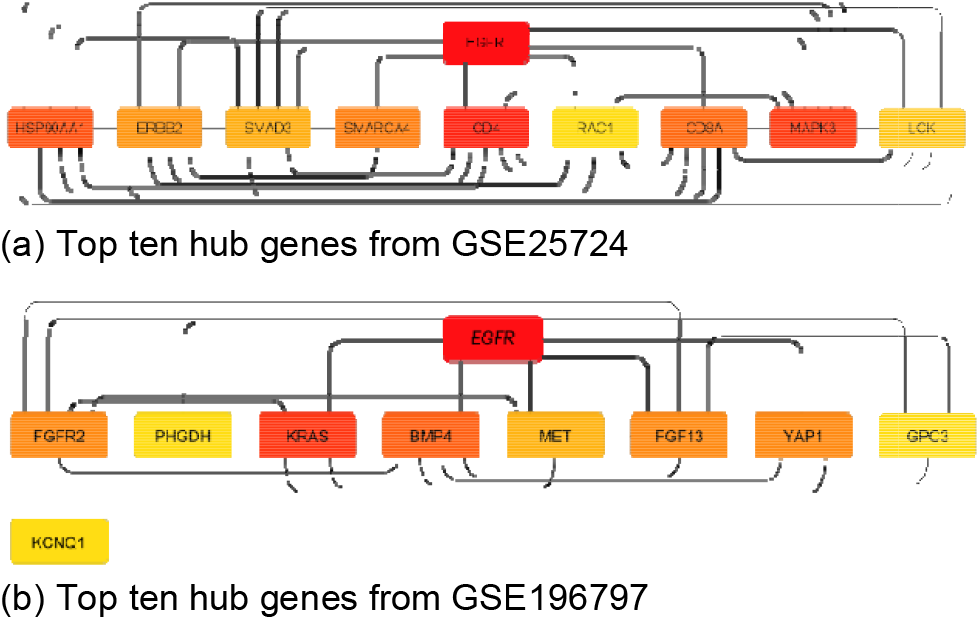
The top ten hub genes with the highest degree of interaction in the PPI network for both studies. Color code for the degree of interaction: darker red for the highest, lighter yellow for the lowest.

**Figure 7.**
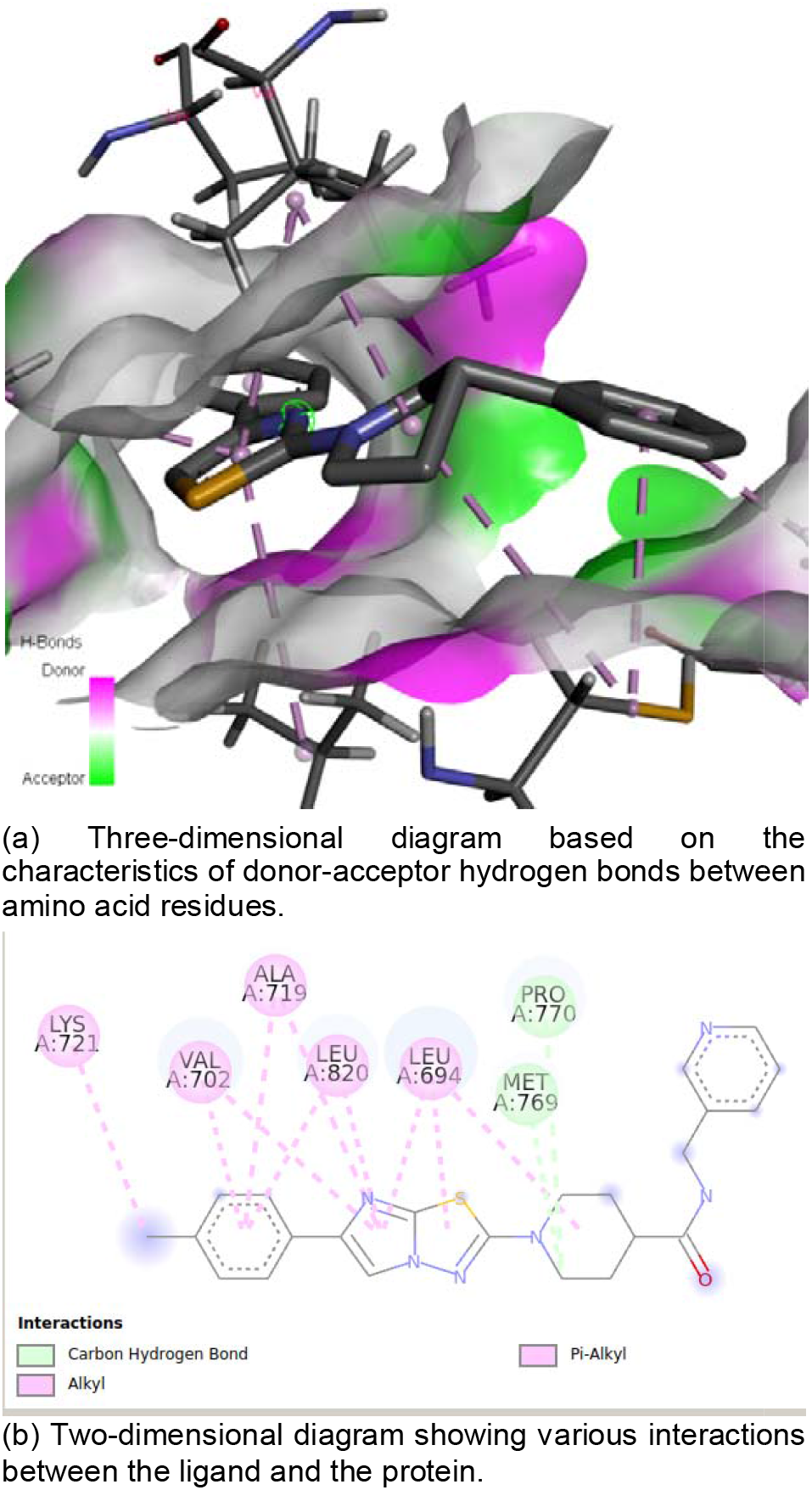
Molecular interactions for the complex EGFR-E626-0366

**Figure 8.**
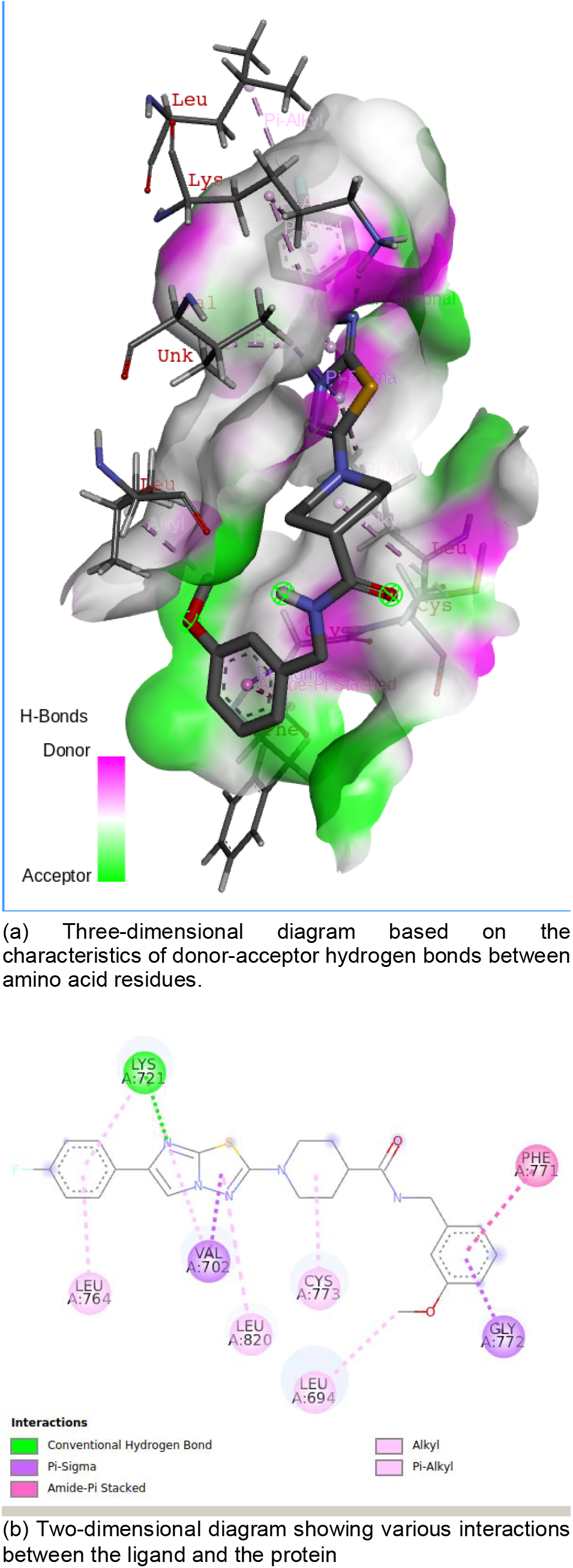
Molecular interactions for the complex EGFR-E626-0262

**Figure 9.**
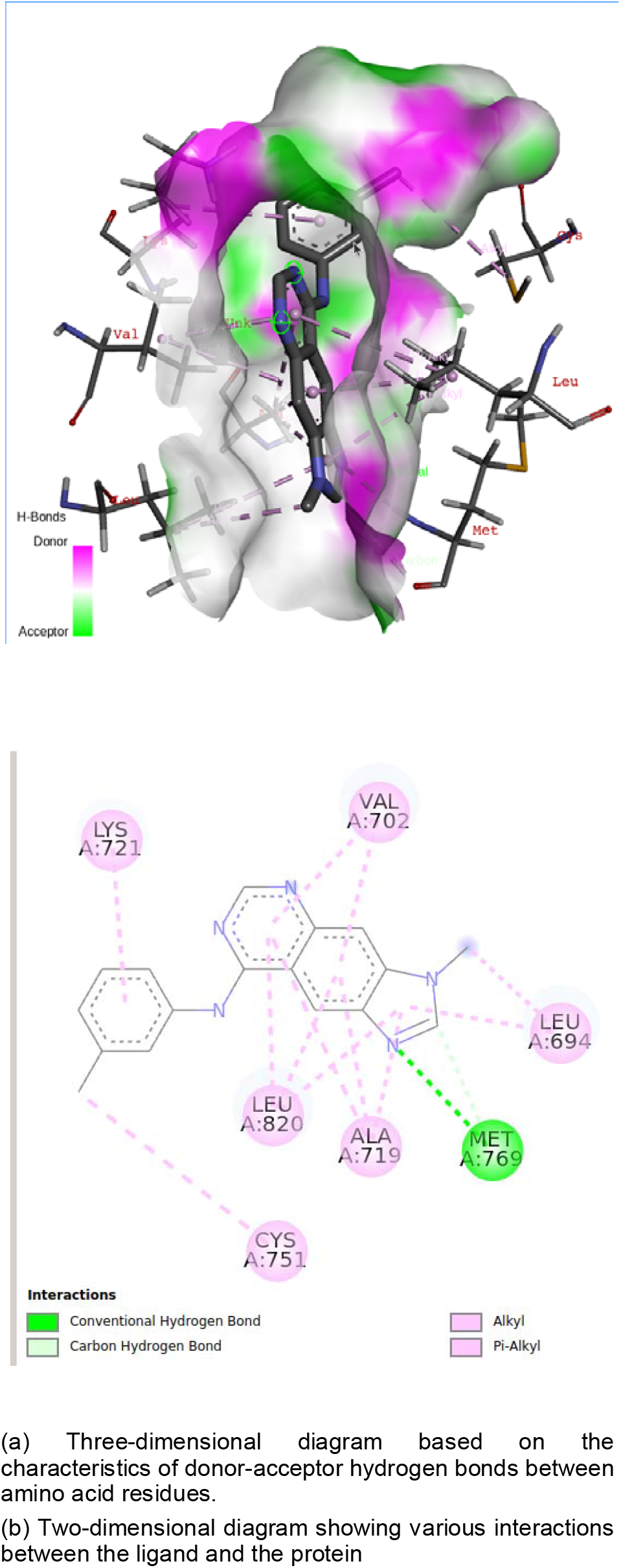
Molecular interactions for the complex EGFR-ZINC6260

Enrichment analysis was performed using hub genes from the GSE25724 study, which demonstrated a superior interaction network (as depicted in Figure 6). The results from Enrichr and GeneTrail are presented in Figure 10. Notably, GeneTrail analysis yielded identical top ten enriched categories regardless of whether the hub genes were directly uploaded, or the GEO accession number was provided. This consistency indirectly verifies the validity of the obtained hub genes. An important finding from the enrichment analysis is the involvement of EGFR in pancreatic cancer signaling pathways. Among the set of hub genes, EGFR, ERBB2, RAC1, MAPK3, and SMAD3 were found to participate in these pathways. The analysis also revealed sets of miRNAs associated with diabetes mellitus and pancreatic neoplasms, which are available in the supplementary materials. Figure 11 depicts the miRNA interaction network.

**Figure 10.**
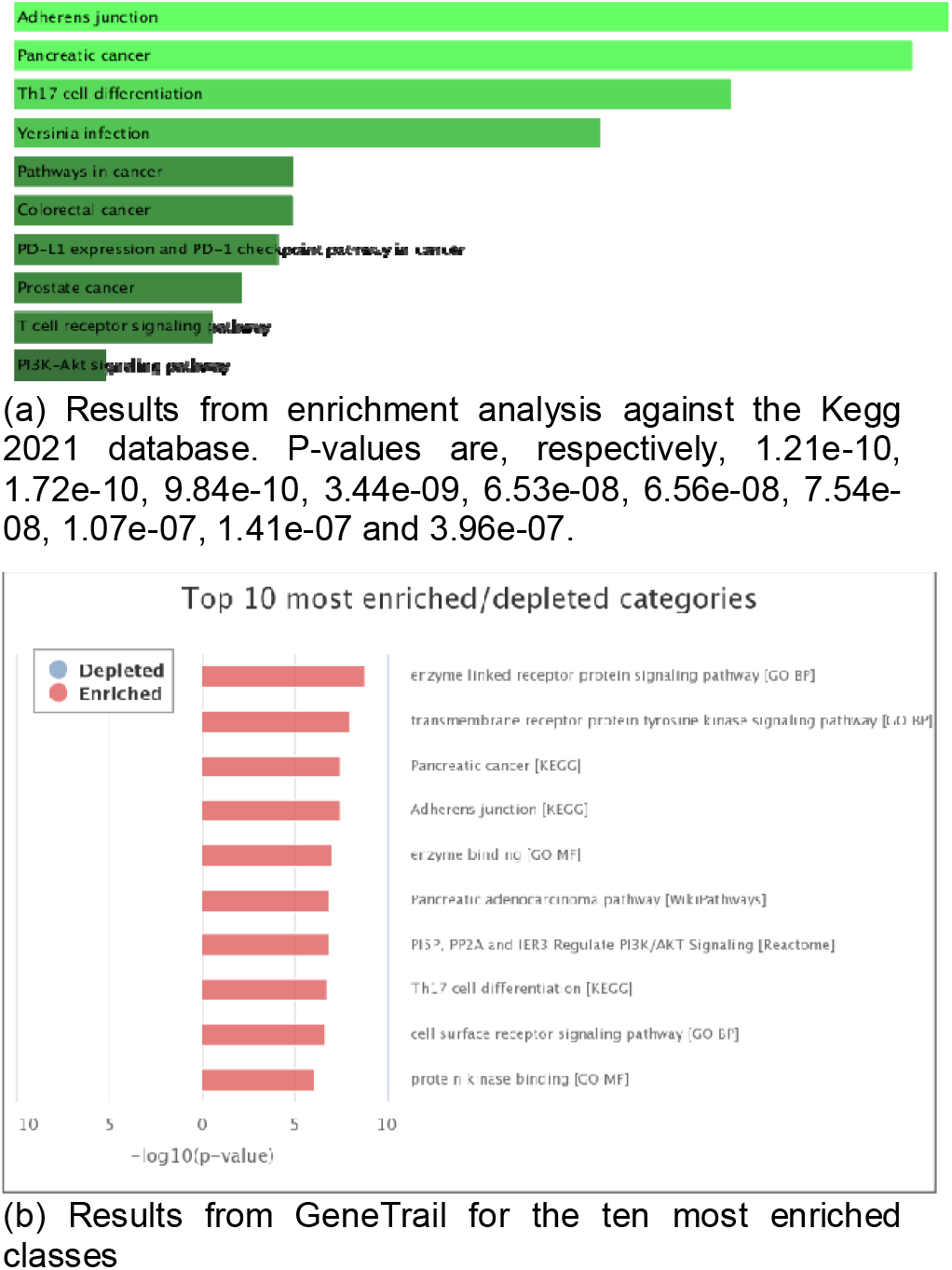
Results from enrichment analysis of the ten hub genes from GSE257

**Figure 11.**
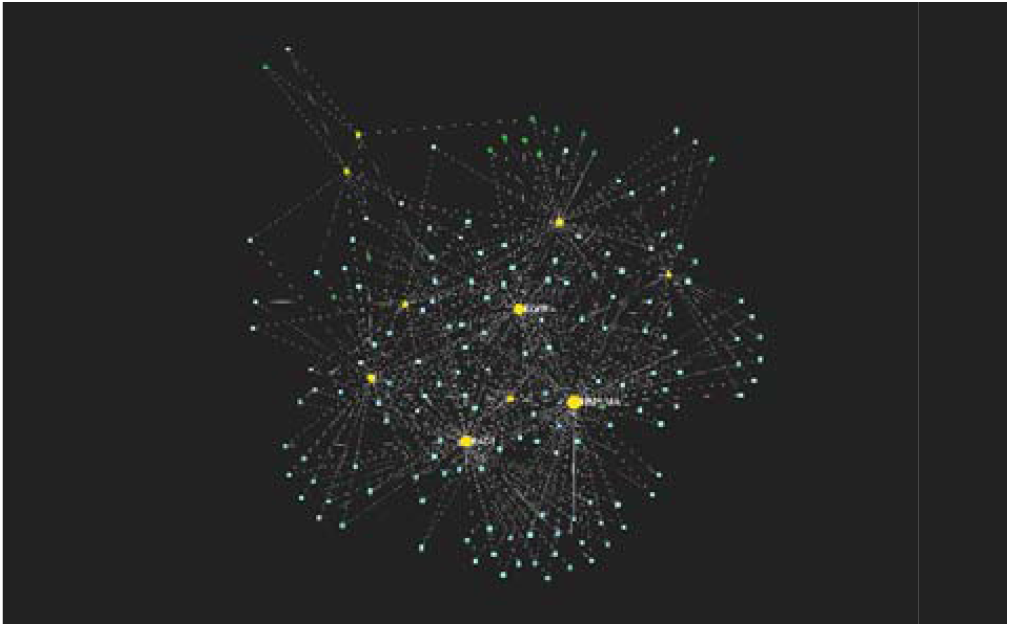
miRNAs (in blue) interaction network obtained from miRNet using the hub genes (in yellow) from the GSE25724 study.

The results of the Mann-Whitney test for active and inactive compounds obtained from ChEMBL, using their Lipinski descriptors (molecular weight, octanol-water partition coefficient, number of hydrogen bond donors and number of hydrogen bond acceptors) as chemical space variables, are presented in Table 5. It can be concluded that the compounds belong to different distributions, but both classes occupy the same chemical space, as shown in Figure 12.

**Table 4.**
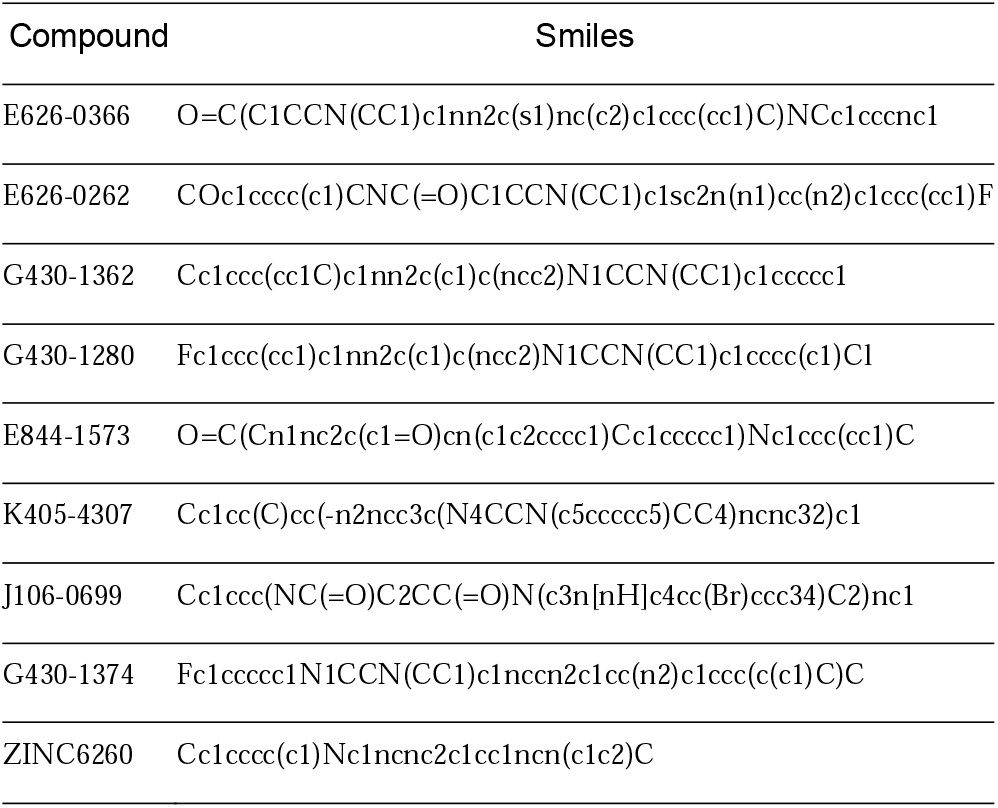
Finalists’ compounds canonical SMILES.

**Table 5.**
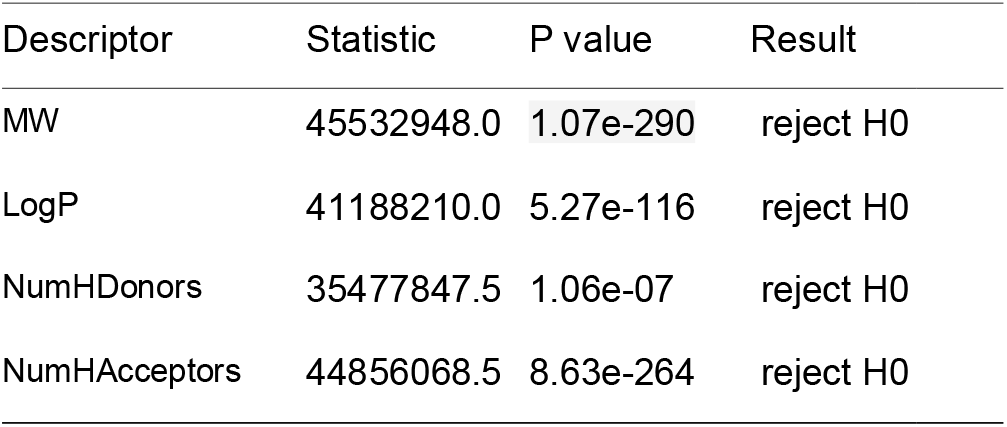
Mann-Whitney test results (α = 0.05) for active and inactive compounds, from ChEMBL, for their Lipinski descriptors: Molecular weight (MW), Octanol – water partition coefficient (LogP), number of Hydrogen bond donors and acceptors. In all cases the null hypothesis is rejected, and we conclude that the compounds belong to different distributions.

**Figure 12.**
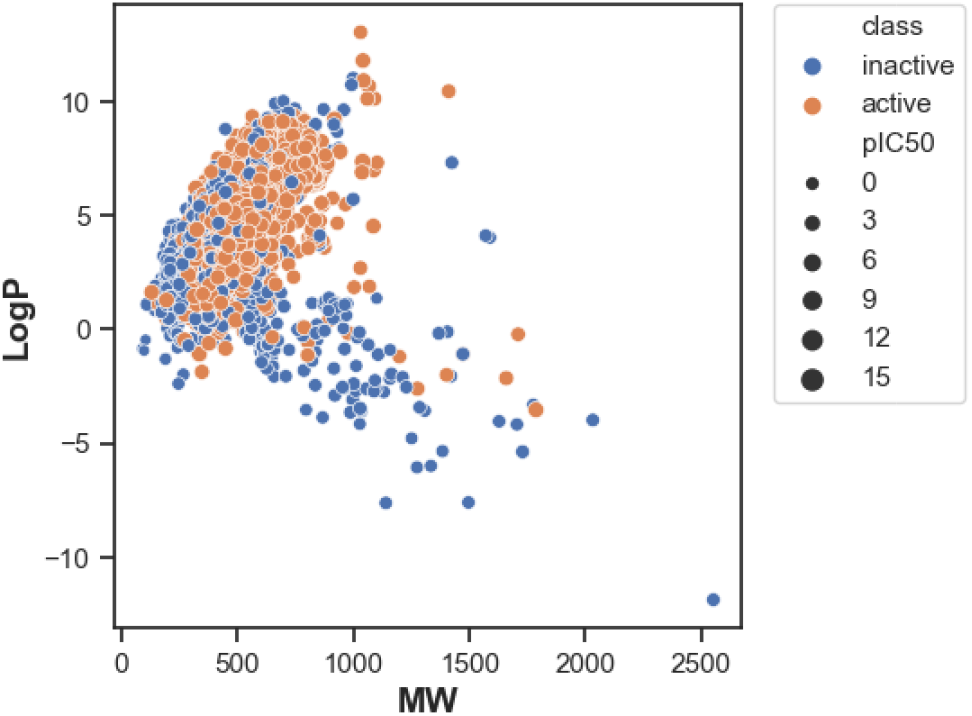
Scatter plot of molecular weight (MW) vs octanol water partition coefficient (LogP) for active and inactive compounds obtained from ChEMBL. The plot shows that both classes occupy the same chemical space despite originating from different distributions based on their Lipinski descriptors.

Lastly, the MLP bioactivity classification model achieved an accuracy rate of 83% for classifying non-active compounds (inactive and intermediate) and an accuracy rate of 86% for classifying active compounds. The average precision rate was 85%, and the AUC value for actual vs predicted probabilities was 0.92. Figure 13 displays the graph and the confusion matrix. As for the Tox21 MLP model it attained a training accuracy of 78% and a test accuracy of 80% showing that it performs well for unseen data. The training loss and validation accuracy scores are plotted in Figure 14.

**Figure 13.**
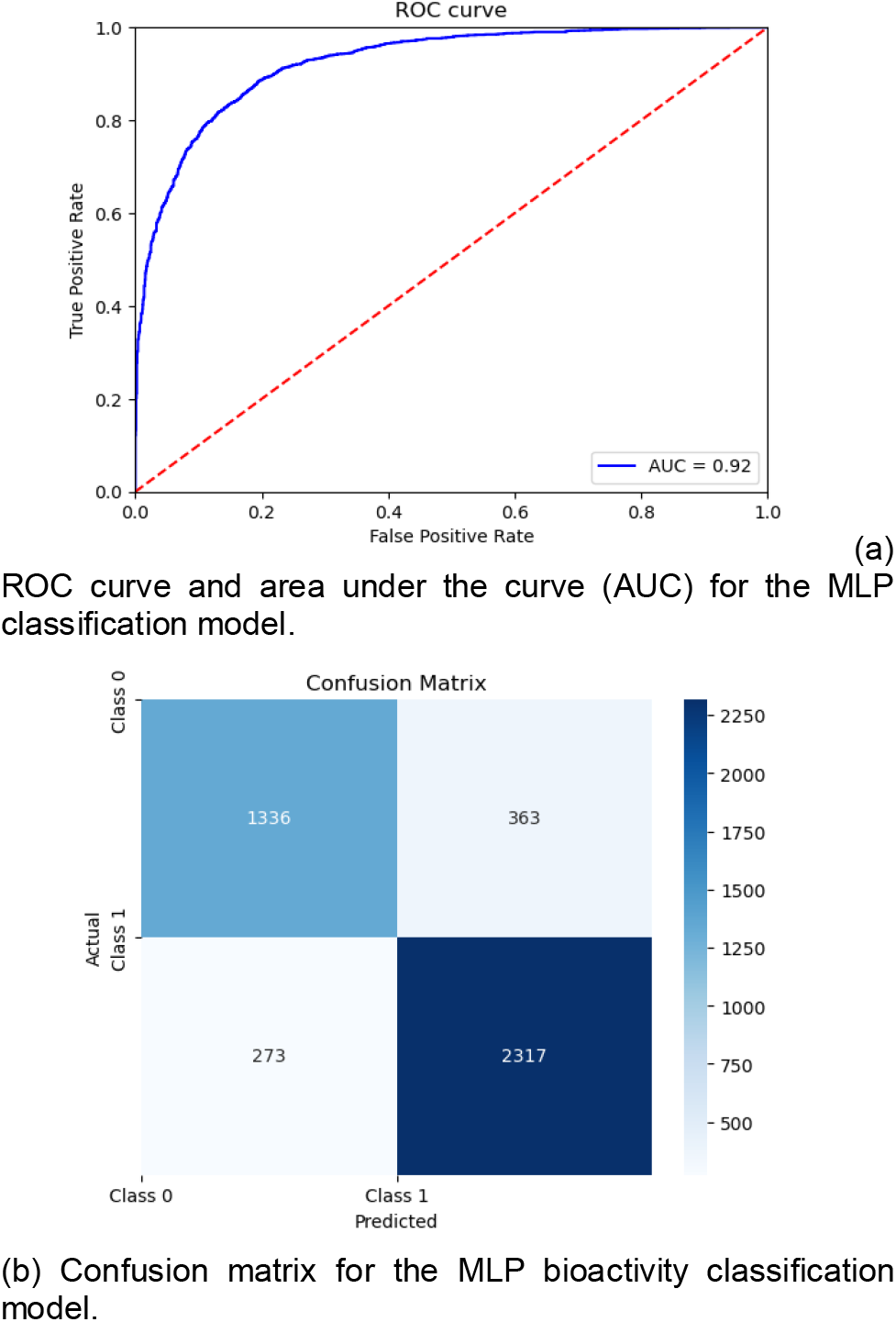
Performance plots for the neural network bioactivity classification model

**Figure 14.**
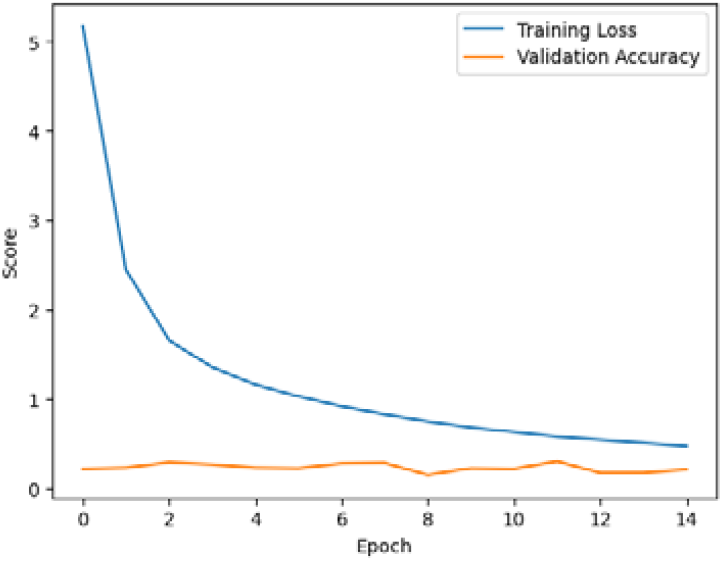
Training loss and validation scores plot for the Tox21 classification model.

## 3 Concluding remarks

The analysis of Tables 1 and 2 demonstrates the superior performance of the finalist compounds compared to the control drugs. The control drugs did not meet the druglikeness and toxicity filters, highlighting the potential limitations of their use. Remarkably, the finalist compounds have not been previously investigated as tyrosine kinase inhibitors, making them novel candidates for further exploration. Based on their scores, the priority for lead development can be determined.

The deep learning models employed in this study have proven to be valuable resources. The availability of both the training data and the models themselves allows for their immediate application in virtual screening of ligands targeting other tyrosine kinase proteins or for training other models. This expands their usability beyond the scope of this work and enhances their research potential.

Another significant finding is the association of the EGFR gene with diabetes, independent of epigenetic studies. While further experimental validation is necessary, our results support existing studies that have reported this association, although it has not been definitively established as mentioned in the introduction. Additionally, the enrichment analysis highlights the involvement of EGFR in the signaling pathway of pancreatic cancer. In summary, these findings strengthen the link between EGFR and type 2 diabetes, and successfully achieve our objective of identifying potential drug candidates for its treatment. The narrowed-down viable hits from this study can now be further investigated in vitro. Furthermore, considering EGFR’s well-known role as an oncogene, these results lay the foundation for the development of improved drugs for diseases requiring EGFR inhibition.

