## Supplementary material for "Identification of Potential EGFR Inhibitors for Type 2 Diabetes and Pancreatic Cancer Treatment: A Computational Approach": Supplemetary material: Diabetes Mellitus miRNAs.pdf

- hsa-let-7a-5p
- hsa-mir-16-5p
- hsa-mir-17-5p
- hsa-mir-21-5p
- hsa-mir-23a-3p
- hsa-mir-30a-5p
- hsa-mir-7-5p
- hsa-mir-34a-5p
- hsa-mir-1-3p
- hsa-mir-133a-3p
- hsa-mir-145-5p
- hsa-mir-9-3p
- hsa-mir-146a-5p
- hsa-mir-185-5p
- hsa-mir-186-5p
- hsa-mir-200a-3p
- hsa-mir-130b-3p
- hsa-mir-30e-5p
- hsa-mir-326
- hsa-mir-196b-5p
- hsa-mir-495-3p
- hsa-mir-550a-3p
- hsa-mir-16-1-3p
- hsa-mir-7-1-3p
- hsa-mir-7-2-3p
- hsa-mir-23b-5p
- hsa-mir-145-3p
- hsa-mir-146a-3p
- hsa-mir-200c-5p
- hsa-mir-342-5p
- hsa-mir-486-3p
- hsa-mir-708-5p
- hsa-mir-133a-5p
- hsa-mir-1-5p
- hsa-mir-130a-3p
- hsa-mir-486-5p
- hsa-mir-9-5p
- hsa-mir-23b-3p
- hsa-mir-103a-2-5p
- hsa-mir-103a-3p
- hsa-mir-107
- hsa-mir-130b-5p
- hsa-mir-181c-5p
- hsa-mir-196a-5p
- hsa-mir-20a-5p
- hsa-mir-21-3p
- hsa-mir-30b-5p
- hsa-mir-30d-5p
- hsa-mir-34a-3p
- hsa-mir-30b-3p
- hsa-mir-708-3p
- hsa-mir-17-3p

- hsa-mir-20a-3p
- hsa-mir-92a-1-5p
- hsa-mir-200b-3p
- hsa-let-7f-5p
- hsa-mir-186-3p
- hsa-mir-196b-3p
- hsa-mir-26b-5p
- hsa-mir-429
- hsa-mir-185-3p
- hsa-mir-16-2-3p
- hsa-mir-200c-3p
- hsa-mir-7

**Pancreatic Neoplasms miRNAs**

- hsa-let-7a-5p
- hsa-let-7b-5p
- hsa-mir-15a-5p
- hsa-mir-17-5p
- hsa-mir-21-5p
- hsa-mir-25-3p
- hsa-mir-27a-3p
- hsa-mir-199a-5p
- hsa-mir-148a-3p
- hsa-mir-30c-5p
- hsa-mir-34a-5p
- hsa-mir-222-3p
- hsa-mir-224-5p
- hsa-mir-15b-5p
- hsa-mir-132-3p
- hsa-mir-145-5p
- hsa-mir-146a-5p
- hsa-mir-186-5p
- hsa-mir-155-5p
- hsa-mir-200a-3p
- hsa-mir-34c-5p
- hsa-mir-133b
- hsa-mir-146b-5p
- hsa-mir-27a-5p
- hsa-mir-29b-1-5p
- hsa-mir-145-3p
- hsa-mir-146a-3p
- hsa-mir-200c-5p
- hsa-mir-155-3p
- hsa-mir-486-3p
- hsa-mir-181b-5p
- hsa-mir-486-5p
- hsa-let-7d-5p
- hsa-let-7g-3p
- hsa-mir-107
- hsa-mir-15a-3p
- hsa-mir-196a-5p
- hsa-mir-20a-5p

- hsa-mir-21-3p
- hsa-mir-24-3p
- hsa-mir-34a-3p
- hsa-let-7c-5p
- hsa-mir-221-5p
- hsa-mir-30c-1-3p
- hsa-mir-30c-2-3p
- hsa-mir-191-5p
- hsa-mir-214-3p
- hsa-mir-17-3p
- hsa-mir-20a-3p
- hsa-mir-96-5p
- hsa-mir-200b-3p
- hsa-let-7e-5p
- hsa-let-7f-5p
- hsa-let-7g-5p
- hsa-let-7i-5p
- hsa-mir-181b-3p
- hsa-mir-186-3p
- hsa-mir-32-3p
- hsa-mir-210-3p
- hsa-mir-106a-5p
- hsa-mir-200c-3p
