## Supplementary material for "Identification of Potential EGFR Inhibitors for Type 2 Diabetes and Pancreatic Cancer Treatment: A Computational Approach": Supplemetary material: Python and R Code.pdf

### ML model for predicting bioactivity

#### Module 1. Data acquisition and cleansing

In [ ]:
```
# Required packages
import pandas as pd
from chembl_webresource_client.new_client import new_client
```

In [ ]:
```
# Search for the target molecule
target = new_client.target
target_search = target.search("egfr")
# Creates a dataframe with the results
target_res = pd.DataFrame.from_dict(target_search)
target_res
```

In [ ]:
```
# Selects the target
target_id = target_res.target_chembl_id[3]
target_id
```

In [ ]:
```
# Adds bioactivity data
activity = new_client.activity
bioactivity = activity.filter(target_chembl_id=target_id).filter(standard_type="IC50")
df = pd.DataFrame.from_dict(bioactivity)
```

In [ ]:
```
# saves results to csv file
df.to_csv("EGFR_raw_data.csv", index=False)
```

In [ ]:
```
# verifies standard_value column (IC50)
# data type
df.standard_value.describe()
```

In [ ]:
```
# converts data to float numbers
df = df.astype({"standard_value":"float"})
df.standard_value.describe()
```

In [ ]:
```
# Removes zero IC50 values
df = df[df.standard_value != 0]
# Removes misssing data
df2 = df.dropna(subset=['standard_value'])
df2 = df2.dropna(subset=['canonical_smiles'])
```

In [ ]:
```
# Removes duplicated canonical smiles
df3 = df2.drop_duplicates(["canonical_smiles"])
```

In [ ]:
```
# Saves preprocessed data
df3.to_csv("EGFR_preprocessed_data.csv")
```

In [ ]:
```
df3 = pd.read_csv('EGFR_preprocessed_data.csv')
```

In [ ]:
```
# Selects relevant columns
select = ["molecule_chembl_id", "canonical_smiles", "standard_value"]
df4 = df3[select]
df4.head()
```

In [ ]:
```
# Creates bioactivity labels
```

```
# inactive >= 10000
# active <= 1000
# rest are intermediate

activity_threshold = []

for i in df4.standard_value:
    if float(i) >= 10000:
        activity_threshold.append("inactive")
    elif float(i) <= 1000:
        activity_threshold.append("active")
    else:
        activity_threshold.append("intermediate")
```

In [ ]:

```
# Creates "class" column
bioactivity_class = pd.Series( activity_threshold, name = "class")
df5 = pd.concat([df4, bioactivity_class], axis = 1)
df5
```

In [ ]:

```
# saves final data
df5.to_csv("EGFR_curated_data.csv", index =False)
```

### ML model for predicting bioactivity

#### Module 2. Statistical analysis

In [ ]:
```
import pandas as pd
import numpy as np   # módulo numérico
from rdkit import Chem  # importa el módulo químico de rdkit
from rdkit.Chem import Lipinski # módulo para calcular descriptores de Lipinski
from rdkit.Chem import Descriptors # módulo para calcular descriptores moleculares
import seaborn as sns # módulo estadístico
sns.set(style="ticks")  # pone marcas en los ejes de las gráficas
import matplotlib.pyplot as plt # módulo gráfico
```

In [ ]:
```
# loads previous data
df = pd.read_csv("EGFR_curated_data.csv")
df.head()
```

In [ ]:
```
# Function to calculate Lipinski descriptors

def lipinski(smiles, verbose=False):

    moldata = []
    for elem in smiles:
        mol = Chem.MolFromSmiles(elem)
        moldata.append(mol)

    baseData = np.arange(1,1)
    i=0
    for mol in moldata:

        desc_MolWt = Descriptors.MolWt(mol)
        desc_MolLogP = Descriptors.MolLogP(mol)
        desc_NumHDonors = Lipinski.NumHDonors(mol)
        desc_NumHAcceptors = Lipinski.NumHAcceptors(mol)

        row = np.array([
            desc_MolWt,
            desc_MolLogP,
            desc_NumHDonors,
            desc_NumHAcceptors
        ])

        if(i==0):
            baseData=row
        else:
            baseData=np.vstack([baseData, row])
        i = i+1

    columnNames=["MW", "LogP", "NumHDonors", "NumHAcceptors"]
    descriptors = pd.DataFrame(data=baseData, columns=columnNames)

    return descriptors
```

In [ ]:
```
df_lipinski = lipinski(df.canonical_smiles)
df_lipinski
```

In [ ]:
```
df2 = pd.concat([df, df_lipinski], axis = 1)
df2
```

In [ ]:
```
# If there are IC50 values equal or greater than 10**9
# they must be normalized to a cut-off value
def norm_value(input):
    norm = []

    for i in input["standard_value"]:
        if i > 10**9:
            i = 10**9
        norm.append(i)

    input["standard_value_norm"] = norm
    x = input.drop("standard_value",1)

    return x
```

In [ ]:
```
# converts IC50 to pIC50 values by applying -log10(IC50)

def pIC50(input):
    pIC50 = []

    for i in input["standard_value_norm"]:
        molar = i*(10**-9)     # convierte los valores de IC50 de nMol a Mol
        pIC50.append(-np.log10(molar)) # convierte de Mol a pIC50

    input["pIC50"] = pIC50
    x = input.drop("standard_value_norm",1)

    return x
```

In [ ]:
```
# applies cut-off and conversion functions
df3 = norm_value(df2)
df4 = pIC50(df2)
df4
```

In [ ]:
```
# saves the final data
df4.to_csv("EGFR_final_data.csv", index=False)
```

In [ ]:
```
# Removes intermediate class
df5 = df4[df4["class"] != "intermediate"]
df5
```

In [ ]:
```
# Compares the two classes

plt.figure(figsize=(3.5,3.5))

sns.countplot(x = "class", data = df5, edgecolor = "black")

plt.xlabel("Bioactivity class", fontsize = 14, fontweight = "bold")
plt.ylabel("Frequency", fontsize = 14, fontweight = "bold")

plt.savefig("bioactivity.png")
```

In [ ]:
```
# Scatter plot of logP vs MW

plt.figure(figsize=(4.5,4.5))

sns.scatterplot(x = "MW", y = "LogP", data = df5, hue = "class", size = "pIC50")
plt.xlabel("MW", fontsize = 14, fontweight = "bold")
plt.ylabel("LogP", fontsize = 14, fontweight = "bold")
plt.legend(bbox_to_anchor = (1.05, 1), loc = 2, borderaxespad = 0)
plt.savefig("MW_vs_LogP.png")
```

In [ ]:
```python
# Box plots for Lipinki descriptors

plt.figure(figsize=(5.5,5.5))

sns.boxplot(x = "class", y = "MW", data = df5)
plt.xlabel("class", fontsize = 14, fontweight = "bold")
plt.ylabel("MW", fontsize = 14, fontweight = "bold")
plt.savefig("Boxplot_MW.png")
```

In [ ]:
```python
plt.figure(figsize=(5.5,5.5))

sns.boxplot(x = "class", y = "LogP", data = df5)
plt.xlabel("class", fontsize = 14, fontweight = "bold")
plt.ylabel("LogP", fontsize = 14, fontweight = "bold")
plt.savefig("Boxplot_LogP.png")
```

In [ ]:
```python
plt.figure(figsize=(5.5,5.5))

sns.boxplot(x = "class", y = "NumHDonors", data = df5)
plt.xlabel("class", fontsize = 14, fontweight = "bold")
plt.ylabel("NumHDonors", fontsize = 14, fontweight = "bold")
plt.savefig("Boxplot_NumHDonors.png")
```

In [ ]:
```python
plt.figure(figsize=(5.5,5.5))

sns.boxplot(x = "class", y = "NumHAcceptors", data = df5)
plt.xlabel("class", fontsize = 14, fontweight = "bold")
plt.ylabel("NumHAcceptors", fontsize = 14, fontweight = "bold")
plt.savefig("Boxplot_NumHAcceptors.png")
```

In [ ]:
```python
# Mann-Whitney test
# The objective is to prove active and inactive classes
# are different distributions according to Lipinski descriptors

def mannwhitney(descriptor, verbose=False):
    from numpy.random import seed
    from numpy.random import randn
    from scipy.stats import mannwhitneyu

    seed(1)

    selection = [descriptor, "class"]
    df = df5[selection]
    active = df[df["class"]== "active"]
    active = active[descriptor]

    selection = [descriptor, "class"]
    df = df5[selection]
    inactive = df[df["class"]== "inactive"]
    inactive = inactive[descriptor]

    stat, p = mannwhitneyu(active, inactive)

    alpha = 0.05
    if p > alpha:
        interpretation = "Equal distributions (accept H0)"
    else:
        interpretation = "Different distributions (reject H0)"

    results = pd.DataFrame({"Descriptor": descriptor,
                            "Statistics": stat,
                            "p":p,
                            "alpha":alpha,
                            "Interpretation": interpretation}, index=[0])
```

```python
    filename = "mannwhitneyu_" + descriptor + ".csv"
    results.to_csv(filename)

    return results
```

In [ ]:
```python
mannwhitney("MW")
```

In [ ]:
```python
mannwhitney("LogP")
```

In [ ]:
```python
mannwhitney("NumHDonors")
```

In [ ]:
```python
mannwhitney("NumHAcceptors")
```

### Toxicity classifier

In [ ]:
```
import pandas as pd
import matplotlib.pyplot as plt
from sklearn.neural_network import MLPClassifier
from sklearn.pipeline import Pipeline
from sklearn.preprocessing import StandardScaler
from sklearn.model_selection import train_test_split, cross_val_score
from rdkit import Chem
from rdkit.Chem import AllChem
from rdkit.ML.Descriptors import MoleculeDescriptors
```

In [ ]:
```
# Load the Tox21 dataset
tox21_data = pd.read_csv('tox21.csv')
tox21_data.head()
```

In [ ]:
```
# Eliminate missing values
tox21_data = tox21_data.dropna()
tox21_data.head()
```

In [ ]:
```
# Define the function to calculate RDKit fingerprints
def calculate_fingerprint(smiles):
    mol = Chem.MolFromSmiles(smiles)
    fp = Chem.RDKFingerprint(mol)
    return fp
```

In [ ]:
```
# Apply the function to the Tox21 dataset
tox21_data['Fingerprint'] = tox21_data['smiles'].apply(calculate_fingerprint)
```

In [ ]:
```
# Split the dataset into features (X) and target (y)
X = list(tox21_data['Fingerprint'])
y = y = tox21_data.iloc[:, 0:12]
```

In [ ]:
```
# Split the data into training and test sets
X_train, X_test, y_train, y_test = train_test_split(X, y, test_size=0.2, random_state=42
```

In [ ]:
```
# Create a pipeline with scaling and MLPClassifier
pipeline = Pipeline([
    ('scaler', StandardScaler()),
    ('mlp', MLPClassifier(hidden_layer_sizes=(100, 50, 25),
                          activation='relu',
                          alpha=0.001,
                          solver='adam',
                          learning_rate='constant',
                          max_iter=200,
                          batch_size='auto',
                          learning_rate_init=0.001,
                          shuffle=True,
                          random_state=42,
                          tol=1e-4,
                          verbose=True,
                          warm_start=False,
                          early_stopping=True,
                          validation_fraction=0.1,
                          n_iter_no_change=10))
])
```

In [ ]:
```
# Fit the model with the Tox21 dataset
pipeline.fit(X_train, y_train)
```

In [ ]:
```
# Evaluate the model on the training set
train_score = pipeline.score(X_train, y_train)
print("Training Accuracy:", train_score)
```

In [ ]:
```
# Evaluate the model on the test set
test_score = pipeline.score(X_test, y_test)
print("Test Accuracy:", test_score)
```

In [ ]:
```
# Set the number of epochs
n_epochs = len(pipeline.named_steps['mlp'].loss_curve_)
```

In [ ]:
```
# Calculate the validation scores
cv_scores = cross_val_score(pipeline, X_train, y_train, cv=n_epochs)
```

In [ ]:
```
# Plot the loss curve
loss_curve = pipeline.named_steps['mlp'].loss_curve_
plt.plot(loss_curve, label='Training Loss')

# Plot the validation curve
plt.plot(range(n_epochs), 1 - cv_scores, label='Validation Accuracy')

plt.xlabel('Epoch')
plt.ylabel('Score')
plt.legend()
plt.show()
```

In [ ]:
```

```

R script for performing DGE análisis

```
# Load required packages
library(DESeq2)
library(GEOquery)

# Set up path and filename for the input data
ex_file <- "gene_exp.csv"

# Load count matrix into DESeq
expression_matrix <- read.csv(ex_file, header = TRUE, row.names = 1)
counts_matrix <- round(expression_matrix)

# Create design matrix specifying experimental treatments.
design_matrix <- data.frame(row.names = colnames(counts_matrix),
                            condition = c(rep("non-diabetic",6),rep("T2D",2)))

# Convert raw counts to normalized expression values using
dds <- DESeqDataSetFromMatrix(countData=counts_matrix,
colData=design_matrix,
                              design=~condition)

# Perform normalization and statistical testing
dds_norm <- DESeq(dds)
```

```
# Perform differential gene expression analysis
dds_dge_res <- results(dds_norm)


# Extract padj, pvalues and log2FoldChange values for all genes
gene_info <- data.frame(padj = dds_dge_res$padj, pvalue =
dds_dge_res$pvalue,
                        log2fc = dds_dge_res$log2FoldChange)


# Save gene_info dataframe to a CSV file
write.csv(gene_info, "GSE196797.csv", row.names=TRUE)
```
